# High Temporal-Resolution Dynamic PET Image Reconstruction Using A New Spatiotemporal Kernel Method

**DOI:** 10.1101/323394

**Authors:** Guobao Wang

## Abstract

Current clinical dynamic PET has an effective temporal resolution of 5-10 seconds, which can be adequate for traditional compartmental modeling but is inadequate for exploiting the benefit of more advanced tracer kinetic modeling. There is a need to improve dynamic PET to allow fine temporal sampling of 1-2 seconds. However, reconstruction of these short-time frames from tomographic data is extremely challenging as the count level of each frame is very low and high noise presents in both spatial and temporal domains. Previously the kernel framework has been developed and demonstrated as a statistically efficient approach to utilizing image prior for low-count PET image reconstruction. Nevertheless, the existing kernel methods mainly explore spatial correlations in the data and only have a limited ability in suppressing temporal noise. In this paper, we propose a new kernel method which extends the previous spatial kernel method to the general spatiotemporal domain. The new kernelized model encodes both spatial and temporal correlations obtained from image prior information and is incorporated into the PET forward projection model to improve the maximum likelihood (ML) image reconstruction. Computer simulations and an application to real patient scan have shown that the proposed approach can achieve effective noise reduction in both spatial and temporal domains and outperform the spatial kernel method and conventional ML reconstruction method for improving high temporal-resolution dynamic PET imaging.

## I. Introduction

Dynamic positron emission tomography (PET) can monitor spatiotemporal distribution of a radiotracer in human body. With tracer kinetic modeling [1], [2], dynamic PET is capable of quantifying physiologically or biochemically important parameters in regions of interest or voxelwise to detect disease status and characterize severity. Traditionally compartmental models are used for kinetic analysis of dynamic PET data [1]. Other advanced tracer kinetic models such as the distributed-parameter model [3] and the adiabatic adapproximations [4] are considered closer to the physiological process than compartmental models. However, those models have not been well explored in dynamic PET because the effective temporal resolution of clinical dynamic PET has been limited to 5-10 seconds with already-compromised image quality. This poor temporal resolution is insufficient to value the use of advanced kinetic modeling in dynamic PET [3].

We aim to improve the effective temporal resolution of clinical dynamic PET imaging to 1-2 seconds. To achieve the high temporal resolution (HTR), short scan durations must be used, which however results in very low counting statistics in the dynamic frames. Image reconstruction from the low-count projection data is extremely challenging because tomography is ill-posed and high noise exists in tomographic measurement of short time frames.

In PET, incorporation of image prior information into image reconstruction has become a popular means to improve the quality of reconstructed images [5], [6]. Prior information can be either local smoothness of neighboring pixels or obtained from a co-registered anatomical MRI image (e.g., [7]–[11]) or CT image (e.g., [12], [13]). Most of existing PET image reconstruction methods employ an explicit regularization form (e.g., the Bowshwer’s approach [14]) to incorporate image priors and can be complex for practical implementation. Regularization-based methods also commonly require a convergent solution to achieve optimal performance, which is computationally costly and can be inefficient for dynamic PET in which many frames need to be reconstructed. Direct reconstruction is another framework which incorporates kinetic modeling into the reconstruction formula [6], [15]–[17]. The method is statistically efficient when the kinetic model type is well known and all voxels in the field of view can be well described using the same kinetic model. However, this assumption is challenging to meet especially for new radiotracers and new clinical applications where the underlying kinetic model can be different from existing models. Any mismatch of kinetic model can induce significant bias propagation in the kinetic images [18].

The recent kernel method [19], [21], [23] encodes image prior information in the forward model of PET image reconstruction and requires no explicit regularization. The kernel method is easier to implement and has been demonstrated more efficient and better improve PET image reconstruction than regularization-based methods [19], [21]. Existing kernel methods mainly explore spatial correlations of image pixels to improve image quality in the spatial domain [19]–[21]. It has been applied to dynamic PET imaging [19], [20], MRI-guided static PET image reconstruction [21], [22], MRI-guided direct PET parametric image reconstruction [23], [24], and optical tomography [25]. These spatial kernels, however, have a limited ability in suppressing noise in the temporal domain. In HTR dynamic PET imaging, noise variation in the temporal domain can be very severe because many short-time frames are used. It is therefore desirable to include temporal prior knowledge in the kernel method to suppress temporal noise.

In this paper, we extend the spatial kernel method to a spatiotemporal kernel method that allows both spatial and temporal correlations to be encoded in the kernel matrix. We propose a separable spatiotemporal kernel to make the method more computationally tractable and easier use. The new spatiotemporal kernel method is expected to achieve substantial noise reduction in the temporal domain in addition to the enhancement on image quality by existing spatial kernels. Note that this new method is different from other ongoing developments that combine the spatial kernel method with direct reconstruction for well-known kinetic models such as the spectral model [23] or the Patlak graphical model [24]. Our focus here is on the reconstruction of HTR time activity curves, allowing new, unknown kinetic modeling to be explored in clinical translation of dynamic PET.

The rest of this paper is organized as follows. We introduce the generalized theory of spatiotemporal kernel method for dynamic PET reconstruction and describe specific kernels in Section II. We then present a computer simulation study in Section III to validate the improvement of the kernel method over existing methods. Section IV presents the result of applying the new method to real patient data of HTR dynamic PET imaging. Finally conclusions are drawn in Section V.

## II. Theory

### A. Dynamic PET Image Reconstruction

For a time frame *m*, we denote the PET image intensity value at pixel *j* by *x*_*j,m*_ and the measurement in detector pair *i* by *y*_*i,m*_. The expectation of the dynamic projection data ***y̅*** = {*y̅*_*i,m*_} is related to the unknown dynamic image *x* = {*x*_*j,m*_} through

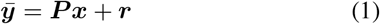

where ***P*** is the detection probability matrix for dynamic PET and *r* is the expectation of dynamic random and scattered events [5].

Dynamic PET projection measurement ***y*** = {*y*_*i,m*_} can be well modeled as independent Poisson random variables with the log-likelihood function [5],

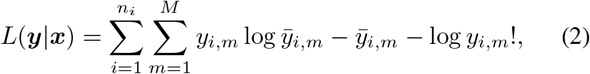

where *n*_*i*_ is the total number of detector pairs and M is the total number of time frames. The maximum likelihood (ML) estimate of the dynamic image *x* is found by maximizing the Poisson log-likelihood,

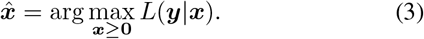

The expectation-maximization (EM) algorithm [26] with the following iterative update

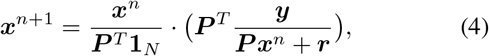

is often the choice to find the solution, where 1_*N*_ is a vector of length *N* = *n*_*i*_ × *M* with all elements being 1, *n* denotes iteration number and the superscript “*T*” denotes matrix transpose. The vector multiplication and division are element-wise operations.

### B. The Spatiotemporal Kernel Method

The kernel method for tomographic image reconstruction [19] was inspired by the kernel methods for classification and regression in machine learning. Different from the kernel methods in machine learning, the kernel method for image reconstruction has unknown “label” values and the available data for kernel coefficient estimation is the tomographic projection data. Previously the kernel method [19] was derived for frame-by-frame spatial image reconstruction, here we adapt the expressions for spatiotemporal reconstruction.

In machine learning language, the image intensity *x*_*j,m*_ at pixel *j* in time frame *m* is the “label” value. For each spatiotemporal location, a set of features are identified to form the feature vector ***f***_*j,m*_, which is also called a “data point” in machine learning. A mapping function *ϕ*(***f***_*j,m*_) is then used to transform the data points {***f***_*j,m*_} into a feature space of very-high dimension {*ϕ*(***f***_*j,m*_}. By doing this, the “label” value *x*_*j,m*_ can be better described as a linear function in the high-dimensional feature space,

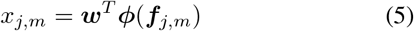

where *w* is a weight vector which also sits in the transformed feature space:

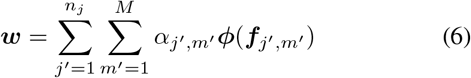

with *α* being the coefficient vector. *n*_*j*_ is the number of pixels in image. By substituting (6) into (5), the kernel representation for the image intensity at pixel *j* and in time frame *m* is written as

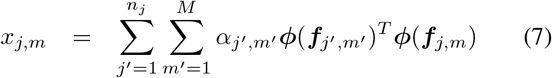

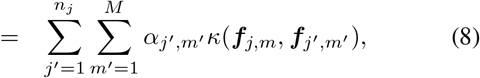

where

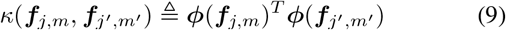

is a kernel defined by the kernel function *κ*(·, ·) (e.g. radial Gaussian function). The mapping function *ϕ* is now implicitly defined by the kernel and not required to be known. The image intensity *x*_*j,m*_ at pixel *j* in time frame *m* is thus described as a linear function in the kernel space but is nonlinear in the original space of the data points {***f***_*j,m*_}. With *x* denoting the dynamic image and ***K*** the spatiotemporal kernel matrix, The equivalent matrix-vector form of (8) is

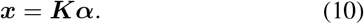

where *α* ≜{*α*_*j,m*_ denotes the kernel coefficient vector.

Substituting the kernelized image model (10) into the standard PET forward projection model (1), we obtain the following kernelized forward projection model for dynamic PET image reconstruction:

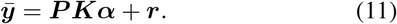

The advantage of using this kernelized model (11) is that image prior knowledge can be incorporated in the forward projection to improve the reconstruction of low-count scans.

A kernelized EM algorithm can be easily derived [19]. The EM update of *α* at iteration (*n* + 1) is

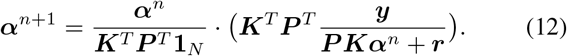

Once the coefficient image *α* is estimated, the reconstructed dynamic PET image is calculated by

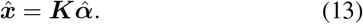

Note that the EM solution converges to the ML estimate when *n* is large, which however can result in noisy reconstruction. In practice, early stopping is commonly used in EM reconstruction to control noise. This mechanism is also used in the kernel-based EM reconstruction.

### C. Separable Spatiotemporal Kernel

The kernel matrix K encodes image prior information based on the feature vectors {***f***_*j,m*_}. For each pixel *j* in time frame *m*, we identify a set of features to form ****f***_*j,m*_,*

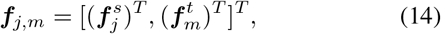

where the vector consists of two components. 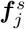 is the vector for exploring spatial correlations between pixels and 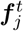 is for exploring temporal correlations between frames.

We further define the spatiotemporal kernel function *κ*(***f***_*j,m*_, ***f***_*j′,m′*_) to be spatially and temporally separable, i.e.

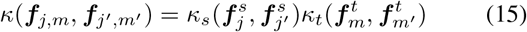

where *κ*_*s*_(·, ·) denotes the kernel function for calculating spatial correlations and *κ*_*t*_(·, ·) is for calculating temporal correlations.

Thus the overall spatiotemporal kernel matrix ***K*** is decoupled into a spatial kernel matrix 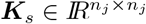 and a temporal kernel matrix 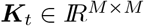,

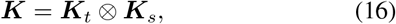

where ⊗ represents the Kronecker product.

Derivation of the spatial kernel matrix ***K***_*s*_ has been developed in our previous work [19]. ***K***_*s*_ is often formed as a sparse matrix based on image prior data. For obtaining image prior in dynamic PET, an effective and efficient means is to use composite frames [19]. For example, an one-hour dynamic FDG-PET scan can be first rebinned into three composite frames, each with 20 minutes. From the reconstructed composite images, three time activity points are obtained at each pixel *j* and used as the feature vector 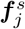 to construct the kernel matrix.

The spatial kernel method [19] is a special example of the spatiotemporal kernel method with the temporal kernel matrix set to the identity matrix,

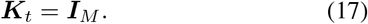

In this paper, we explore the role of new ***K***_*t*_ in the context of HTR dynamic PET imaging.

### D. Choice of Temporal Kernels

The (*m*, *m′*)th element of ***K***_*t*_ is obtained by comparing the feature vectors of the frames *m* and *m′*:

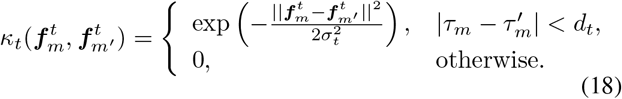

where *τ*_*m*_ denotes the middle time point of frame *m* and *d*_*t*_ is the width of time window for neighborhood construction. *σ*_*t*_ is a parameter to adjust the weight calculation.

The simplest form of the temporal feature 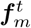 is probably

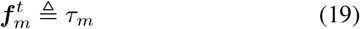

by which ***K***_*t*_ becomes a shift-invariant Gaussian smoothing kernel with the parameter *σ*_*t*_ determined by the window size This type of kernel can smooth out noise but may also over smooth sharp signals in the temporal domain.

To make the temporal kernel more adaptive to time varying data, we propose to use the whole sinogram of each frame as the feature vector to capture temporal correlations between frames, i.e.

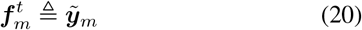

where ***ỹ***_*m*_ denotes a smoothed version of the raw sinogram ***y***_m_ of frame *m*. The parameter *σ*_*t*_ can be set to the standard deviation of all elements of 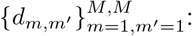

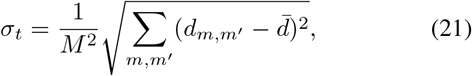

where *d̄* is the mean of {*d*_*m*,*m′*_} and

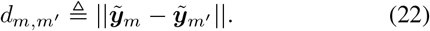

As both ***K***_*s*_ and ***K***_*t*_ are very sparse, inclusion of the kernel matrix ***K*** in the projection model does not add a significant computational cost in the reconstruction.

## III. Validation Using Computer Simulation

### A. Simulation Setup

Dynamic ^18^F-FDG PET scans were simulated for a GE DST whole-body PET scanner in two-dimensional mode using a Zubal head phantom [Figure 1(a)]. The phantom contains several brain regions including brain background, blood region, gray matter, white matter and a tumor (15 mm in diameter). An attenuation map was simulated with a constant linear attenuation coefficient assigned in the whole brain. The scanning schedule consisted of 63 time frames over 20 minutes: 30×2 s, 12×5 s, 6×30 s, 15×60 s.

The blood input function was extracted from a real patient ^18^F-FDG PET scan [Figure 1(b)]. Regional time activity curves (TACs) [Figure 1(c-d)] were assigned to different brain regions to generate noise-free dynamic activity images. The early phase of these TACs has fast temporal dynamics and is challenging to reconstruct. The resulting noise-free activity images were first forward projected to generate noise-free sinograms. A 20% uniform background was included to simulate random and scattered events. Poisson noise was then generated with 20 million expected events over 20 minutes.

**Fig. 1.**
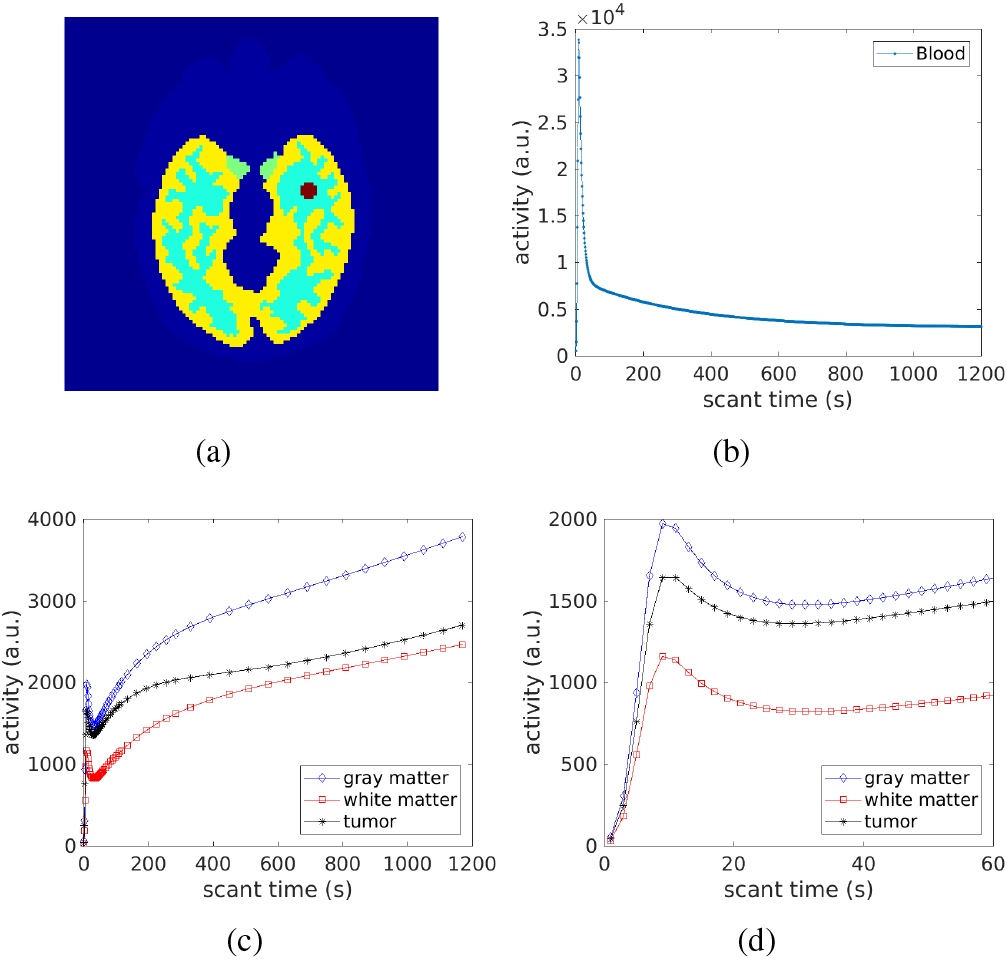
The digital phantom and time activity curves used in the simulation study. (a) The Zubal brain phantom; (b) Blood input function; (c) Regional time activity curves of different brain regions; (d) Time activity curves of the first 60 seconds.

A total of 20 realizations were simulated and each was reconstructed independently for statistical comparison. As the main goal is to compare the reconstruction methods for HTR dynamic PET imaging, the study focused on the comparisons for the first one minute (thirty 2-second frames) of the dynamic scan.

### B. Illustration of Temporal Kernels

One way of understanding the temporal kernel matrix ***K***_*t*_ is that each column of ***K***_*t*_ represents a temporal basis function. For example, the *m*th column of a ***K***_*t*_ represents the temporal basis function centered at the *m*th time frame. Thus, a linear combination of these temporal basis functions consists in a time activity curve.

For illustration, a temporal window size of 15 time frames was used to construct temporal kernels. This fixed number of frames for defining the temporal window leads to varying time windows across the dynamic sequence. The time window size was 30 seconds for early time frames and 15 minutes for late time frames. For calculating the data-driven temporal kernel matrix, the sinogram of each frame was smoothed using Gaussian smoothing with a window size of 7 × 7 before it was used.

Fig. 2 shows the comparison of temporal basis functions of two types of kernels - Gaussian temporal smoothing kernel (Eq. 19) and data-driven temporal kernel (Eq. 20). These temporal bases correspond to the 5th frame (t=10-s), 15th frame (t=30-s) and 55th frame (t = 12-minute), respectively. For early time frames where fast activity changes occur, the data-driven kernel provides sharper temporal basis functions than the Gaussian smoothing kernel, while the two kernels are similar for late-time frames due to the slower change of activities. Thus we expect the data-driven kernel to improve image reconstruction mainly for early-time frames when compared with the Gaussian temporal kernel.

**Fig. 2.**
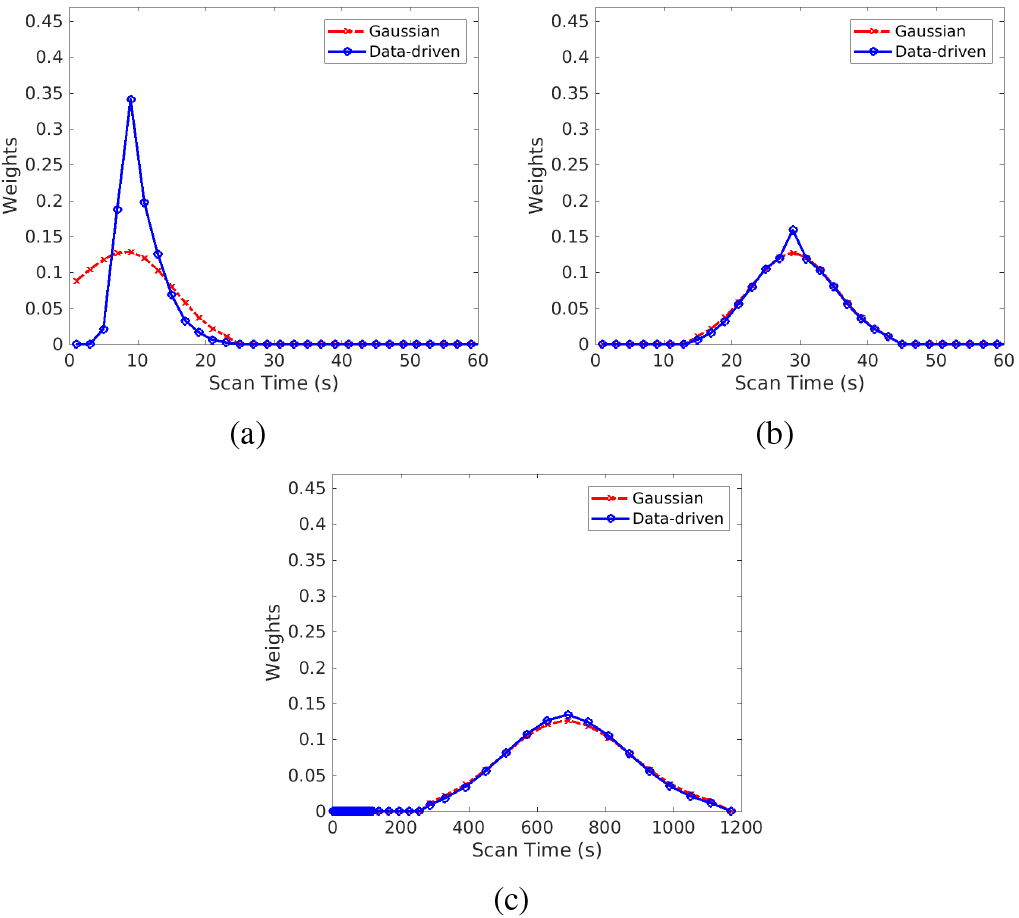
Temporal basis functions formed by the Gaussian kernel and data-driven kernel. (a) center located at 5th frame (*t* = 10 s); (b) centered at 15th frame (*t* = 30 s); (c) centered at 55th frame (*t* = 12 minutes)

Note that the spatial kernel method has a special temporal kernel matrix - the identity matrix ***I***_*M*_, of which the temporal bases correspond to unit impulse functions for all frames. Because there are no temporal correlations included in this impulse kernel, temporal noise will substantially remain in the reconstruction.

### C. Reconstruction Methods to Compare

Noisy sinograms were reconstructed independently by four different image reconstruction methods: the traditional MLEM method, spatial kernelized EM (KEM-S), and new spatiotemporal kernel method with the Gaussian smoothing temporal kernel (KEM-ST-G) and the spatiotemporal kernel method with the data-driven temporal kernel (KEM-ST-D). Each reconstruction was run for 200 iterations to allow the investigation of the effect of iteration number.

The spatial kernel matrix ***K***_*s*_ was constructed using the k nearest neighbor (kNN) approach described in [19]. Four composite images (4 × 5-minute) were obtained by recombining the full 20-minute dynamic data and used for the spatial kernel matrix construction. The temporal kernels were constructed using a window size of 15 frames unless specified otherwise. Different window sizes, ranging from 7 to 33 frames, were also investigated to evaluate the effect of temporal window size on reconstruction quality.

**Fig. 3.**
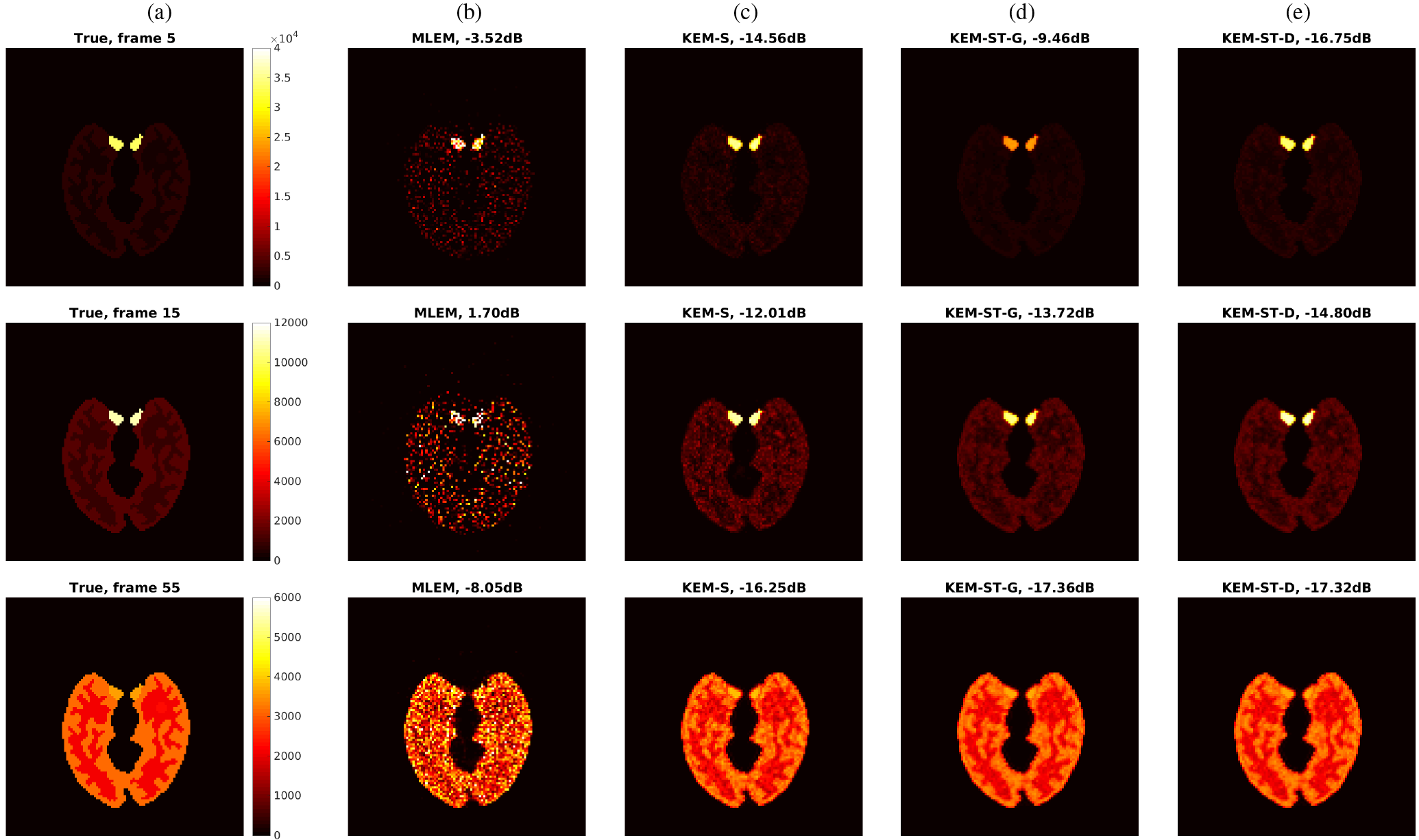
True activity image and reconstructed images at iteration 100 by different reconstruction methods for the 5th frame (*t* = 10 s, top row), 15th frame (*t* = 30 s, middle row) and 55th frame (*t* = 12 minutes, bottom row). (a) True images, (b) MLEM reconstructions, (c) reconstructions by the spatial kernel method (KEM-S), (c) by the spatiotemporal kernel method with the time-invariant Gaussian smooth kernel (KEM-ST-G), (e) by the spatiotemporal kernel method with a data-driven temporal kernel (KEM-ST-D).

**Fig. 4.**
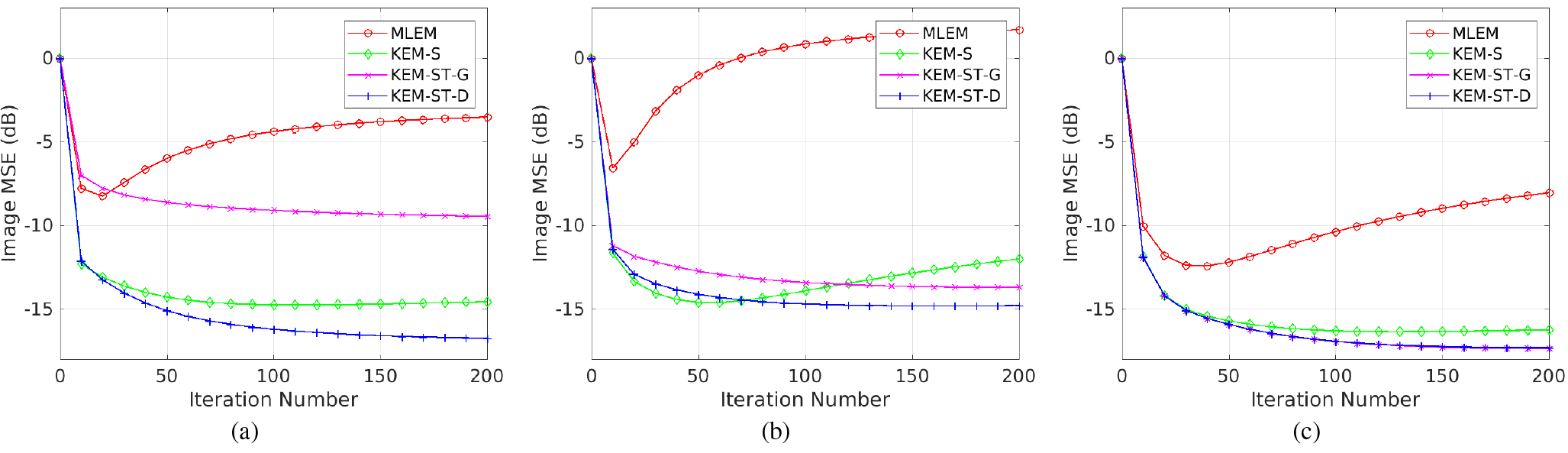
Plots of image MSE as a function of iteration number for different reconstruction methods. (a) frame 5, (b) frame 15, (c) frame 55.

### D. Comparison for Overall Image Quality

Image quality of different reconstruction methods were first assessed using the image mean squared error (MSE) which is defined by

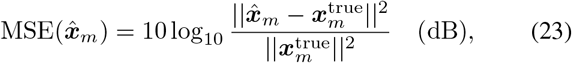

where *x̂*_*m*_ is an image estimate of frame *m* obtained with one of the reconstruction methods and 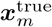 denotes the ground truth image of the frame.

Figure 3 shows the true activity image and reconstructed images by the four different methods with 100 iterations for the 5th frame (*t* = 10 s, Δt = 2 s), 15th frame (*t* = 30 s, Δt = 2 s) and 55th frame (*t* = 12 minutes, Δt = 60 s), respectively. As expected, the KEM-S method achieved a significant MSE decrease as compared with the MLEM method. By incorporating temporal correlations, the KEM-ST-G and KEM-ST-D methods further improved the image quality and achieved lower MSE values than KEM-S. The KEM-ST-D method had lower MSE than the KEM-ST-G method for the early-time frame 5 and frame 15. The two methods had similar MSE for the late-time frame 55.

Figure 4 shows image MSE as a function of iteration number for the three different frames (5th, 15th and 55th). The two spatiotemporal kernel methods KEM-ST-G and KEM-ST-D had slower convenience than the spatial kernel method KEM-S, requiring more iterations to achieve the lowest MSE among iterations. This is because of the additional temporal correlation included in the spatiotemporal kernel methods. Note that the MSE curves of the 15th frame (*t* = 30 s) behaved slightly different from that of the 5th and 55th frames and achieved the lowest MSE earlier. This is because the 15th frame had lower activity and count level than the other two frames [see Fig.1 (c-d) for a comparison].

**Fig. 5.**
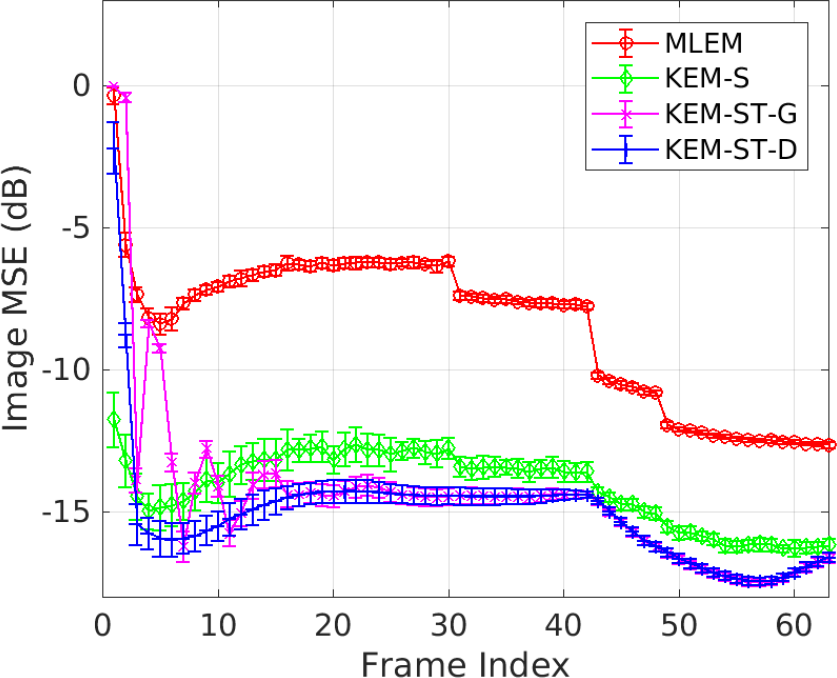
Plots of image MSE of all time frames reconstructed by different reconstruction methods.

Figure 5 shows the plots of image MSE of all time frames for different methods. The MSE of each frame is minimized over iteration numbers in different methods. The error bars indicate the standard deviation of the MSE over the 20 realizations. The three kernel methods outperformed the MLEM reconstruction for all frames. Compared with the spatial kernel method KEM-S, the spatiotemporal kernel method KEM-ST-G improved late-time frames. This is because the TACs of these late-time frames have small temporal changes than early-time frames. Incorporation of temporal correlations thus became beneficial and achieved noise reduction. KEM-ST-D and KEM-ST-G had similar performance given the two temporal kernels were close to each other in the late-time frames. In the early-time frames where activity change is fast, use of the time invariant temporal kernel in the KEM-ST-G method over-smoothed the temporal signals. Thus KEM-ST-G had even higher MSE than KEM-S. In contrast, KEM-ST-D had data-adaptive temporal kernels and achieved better results.

### E. Comparison for Time Activity Curves

Figure 6(a) and 6(b) respectively show the TACs of a pixel in the blood region and a pixel in the tumor region reconstructed by different methods. The TACs of the first 60 seconds are further shown in Figure 6(c) and 6(d). The reconstructions by MLEM were extremely noisy, especially in the early-time frames because these frames are of only 2-s scan duration. While it achieved a significant noise reduction in the spatial domain, the KEM-S method still resulted in substantial noise in the temporal domain, particularly in the tumor where the tracer uptake is relatively low. The KEMST-G method reduced temporal noise in late-time frames but over-smoothed the early-time peak activities, causing bias in early-time frames. In comparison, the KEM-ST-D reconstruction overcame the limitation of KEM-ST-G and achieved a substantial noise reduction in the temporal domain for both early-time and late-time frames.

To compare different methods quantitatively, we calculated the bias and standard deviation (SD) of the mean uptake in the tumor region by

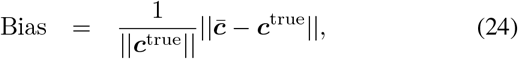

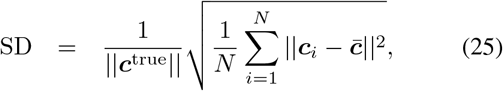

where *c*^true^ is the noise-free regional TAC and 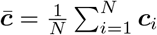 denotes the mean of *N* realizations. *N* = 20 in this study.

Figure 7 shows the trade-off between the bias and SD of different methods in the tumor ROI for two single time points (frame 5 and frame 15) and for the whole TAC of the first 60 seconds. The curves were obtained by varying the iteration number from 10 to 200 iterations with an interval of 10 iterations. Compared with the MLEM and KEM-S methods, the two spatiotemporal kernel methods (KEM-ST-G and KEM-ST-D) had lower noise SD thanks to the temporal kernels. Due to being more adaptive to the data, the KEM-ST-D method achieved lower bias than the KEM-ST-G method at any level of noise.

### F. Effect of Temporal Window Size

Figure 8 shows the MSE of regional TAC of the blood region and tumor region as a function of temporal window size. With the window size equal to 1, the spatiotemporal kernel methods KEM-ST-G and KEM-ST-D become the same as the spatial kernel method KEM-ST.

Because the blood region has a high count level (due to high activity), no temporal smoothing was needed. With increasing temporal window size, the Gaussian kernel ST-G reduced the TAC quality due to over-smoothing. The data-driven kernel ST-D was less affected by the window size and was more stable to maintain the blood TAC. In the tumor region, the performance of the ST-G kernel changed sharply, indicating it can be difficult to choose a proper window size. In contrast, the performance of the ST-D kernel was stable for a range of window sizes from 10 to 30 frames. The minimum MSE achieved by the ST-D kernel was also lower than that by the ST-G kernel.

These results indicate that the ST-D kernel is more adaptive to the data and more stable than the ST-G kernel.

## IV. Application to Real Patient Scan

### A. Patient Data Acquisition

A cardiac patient scan was performed on the GE Discovery ST PET/CT scanner at the UC Davis Medical Center in twodimensional mode. The patient received 20 mCi ^18^F-FDG with a bolus injection. Data acquisition commenced right after the FDG injection. A low-dose transmission CT scan was then performed at the end of PET scan to provide CT image for PET attenuation correction. The raw data of first 90 seconds were binned into a total of 45 dynamic frames, each with 2-second duration. The data correction sinograms of each frame, including normalization, attenuation correction, scattered correction and randoms correction, were extracted using the vendor software used in the reconstruction process.

**Fig. 6.**
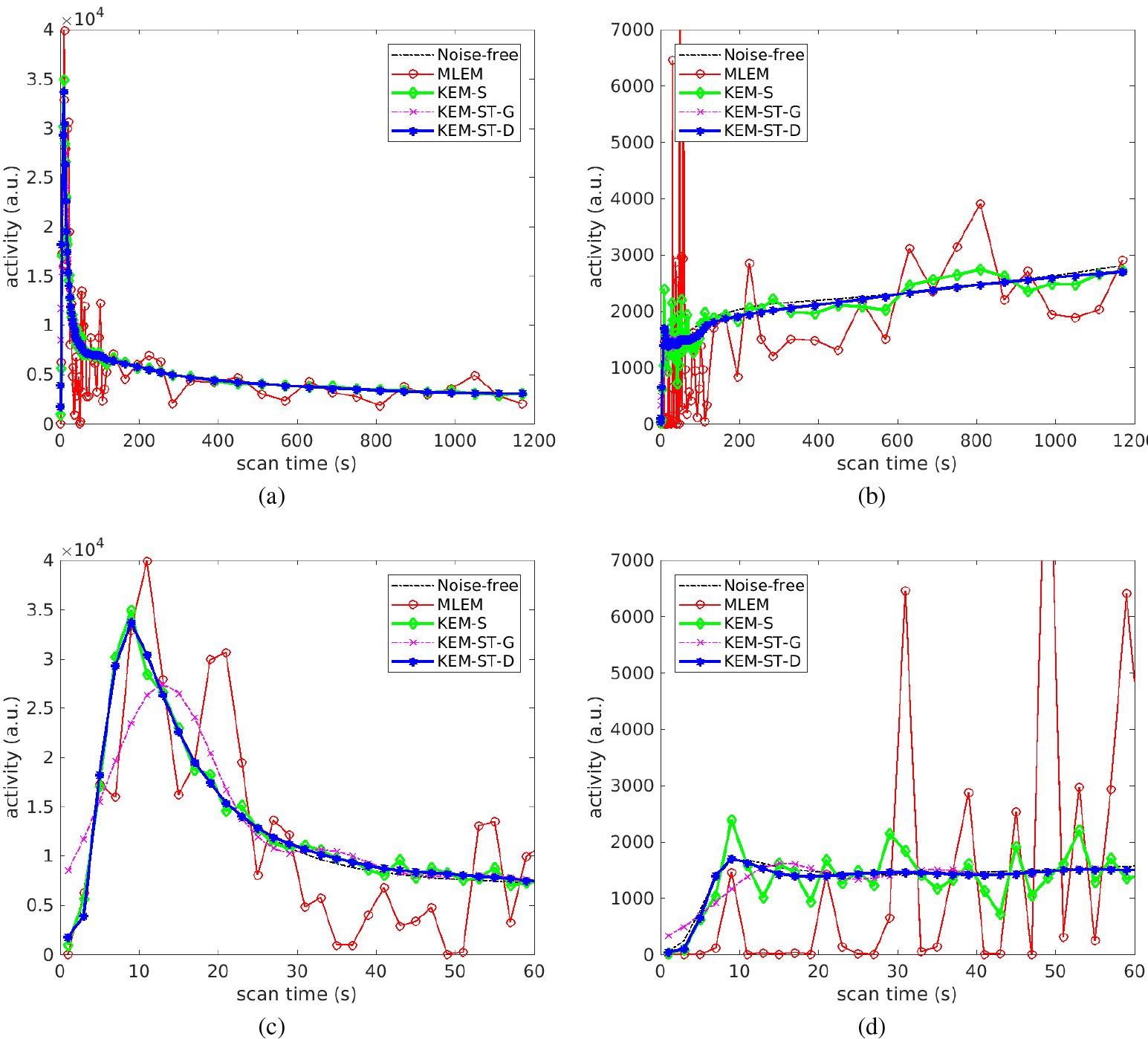
Time activity curves reconstructed by different reconstruction methods. (a) Blood TAC, 0-20 minutes, (b) tumor TAC, 0-20 minutes, (c) blood TAC, 0-60 seconds, (d) tumor TAC, 0-60 seconds.

**Fig. 7.**
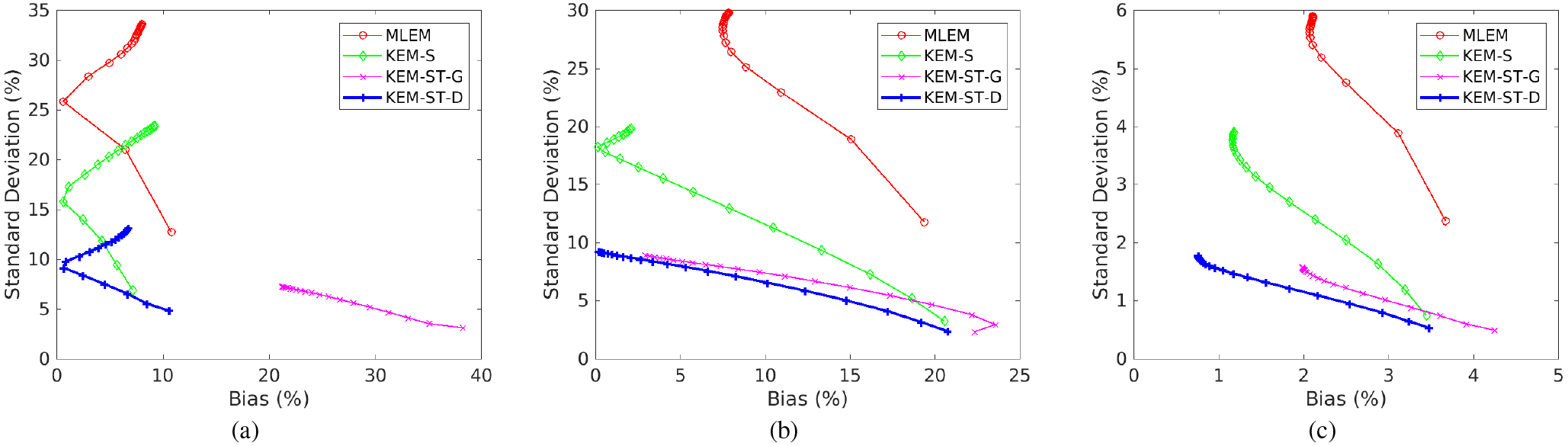
Standard deviation versus bias trade-off of tumor ROI quantification by varying iteration numbers from 10 to 200 iterations. (a) frame 5 (*t* = 10 s), (b) frame 15 (*t* = 30 s), (c) whole TAC of the first 60 seconds.

The patient data were reconstructed independently by the traditional MLEM method and three kernel emthods: spatial kernel method KEM-S, new spatiotemporal kernel methods KEM-ST-G (Gaussian smoothing temporal kernel) and KEM-ST-D (data-driven temporal kernel) with 100 iterations. The spatial kernel matrix ***K***_*s*_ was constructed using five composite frames (3×10 s, 2×30 s). The temporal kernels were constructed using the same parameters as used in the simulation study.

**Fig. 8.**
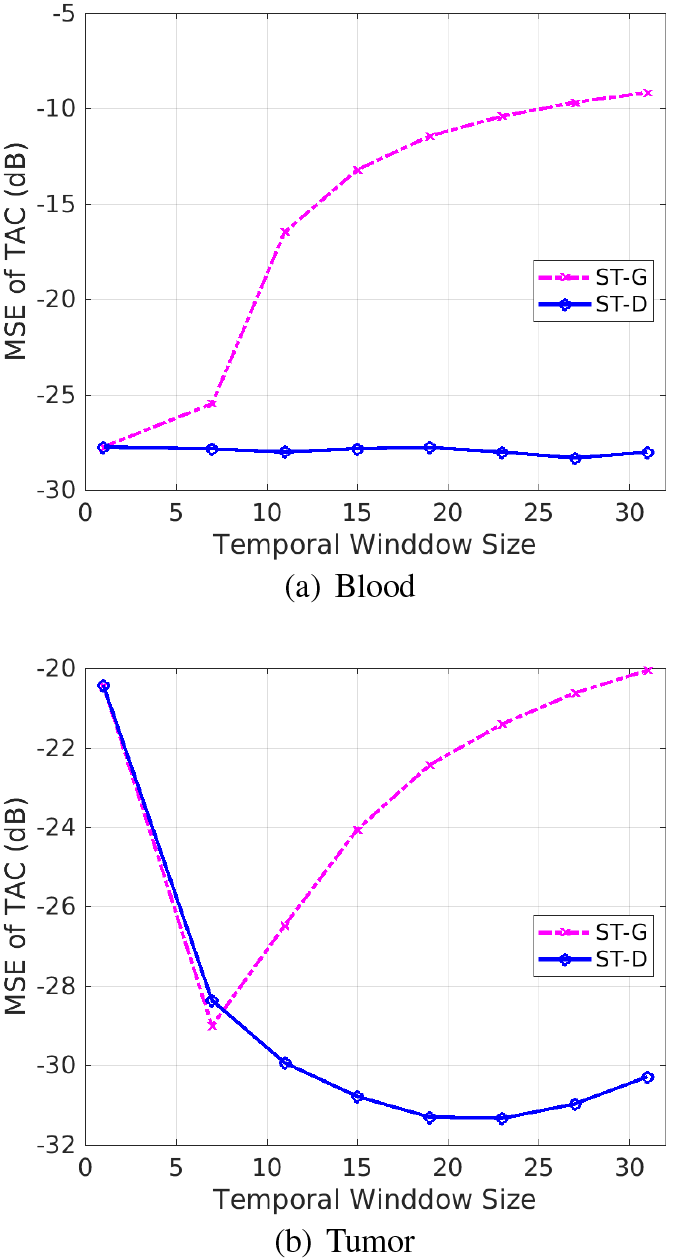
Effect of temporal window size on quantification of regional TACs in the simulation study. (a) Blood ROI, (b) Tumor ROI.

### B. Results

Figure 9 shows the comparison of a 10-s low temporal-resolution time frame by standard MLEM with its corresponding 2-s high temporal-resolution time frames reconstructed using the data-driven spatiotemporal kernel method KEM-ST-D. While the 10-s image by MLEM already suffered from noise and showed similar FDG uptakes in the left ventricle and right ventricle, the HTR images demonstrated time-changing uptakes in the two ventricular regions with good image quality even if they are only of 2-s short duration. The right ventricle had decreasing activities and the left ventricle had increasing activities during the same 10 seconds period.

Figure 10 shows the comparison of different image reconstruction methods for reconstructing HTR time frames at *t* = 10s, *t* = 30s and *t* = 90s. The traditional MLEM reconstructions were extremely noisy. The spatial kernel method KEM-S achieved substantial noise reduction. Compared with KEM-S, KEM-ST-G underestimated the FDG uptake in the left and right ventricles due to Gaussian temporal smoothing. Thanks to being more adaptive to the data than KEM-ST-G, KEM-ST-D preserved the improvement by KEM-S in the early frames at *t* = 10s and *t* = 20s and further reduced the noise in the late frame at *t* = 90s.

Figure 11 shows the HTR time activity curves for the pixels in the right ventricle, left ventricle, aorta and myocardium, respectively. The time activity curves of standard temporal resolution by MLEM are also included for comparison. With the standard MLEM reconstruction, the increase of temporal resolution suffered high noise, thus contaminating any benefit brought by HTR. The kernel-based reconstruction by KEM-S demonstrated low noise for the high-activity time points. However, high noise remained in the low-activity time points. KEM-ST-G smoothed out the noise in late frames but over-smoothed early frames where activity changes fast. In comparison, the KEM-ST-D method achieved satisfied noise suppression for both early-time and late-time points on the time activity curves.

**Fig. 9.**
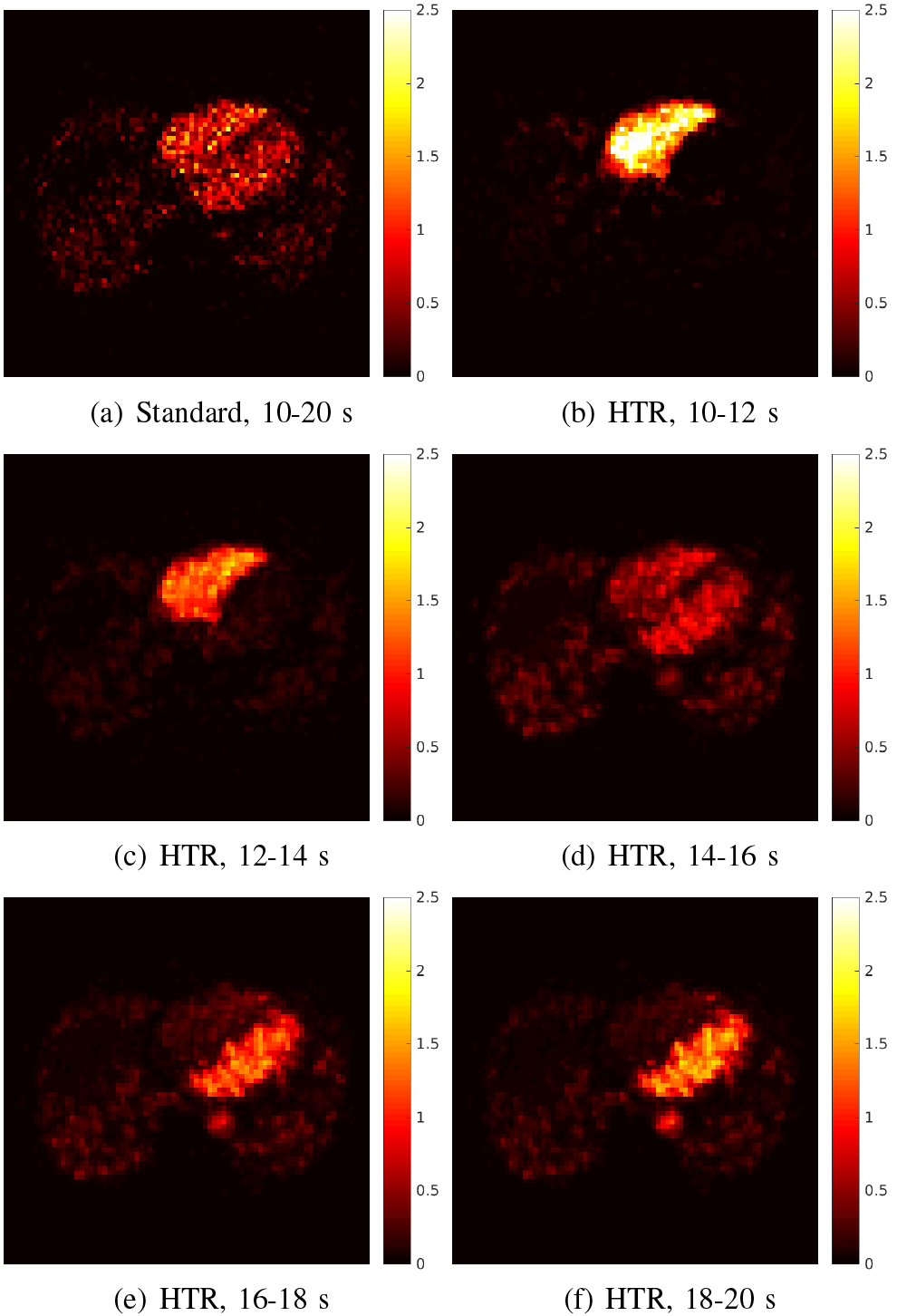
Comparison of a low-resolution time frame starting from 10 to 20s with its corresponding 2-s HTR time frames by the spatiotemporal kernel method for the patient study. (a) 10-s frame at 10-20s by MLEM, (b-f) HTR time frames by KEM-ST-D: (b) 10-12s, (c) 12-14s, (d) 14-16s, (e) 16-18s, (f) 18-20s.

These patient results have further demonstrated the improvement of the spatiotemporal kernel method for HTR image reconstruction in addition to the results from the simulation study.

## V. Conclusion

In this paper, we have developed a spatiotemporal kernel method to incorporate both spatial and temporal prior information into the kernel framework for dynamic PET image reconstruction. The spatiotemporal kernel is separable in the spatial and temporal domains and thus can be easily and efficiently implemented. We conducted a computer simulation to validate the method and tested the method using a patient PET scan. The results from both the simulation study and patient study have shown that the new spatiotemporal kernel method can outperform traditional MLEM and existing spatial kernel methods and achieve a 2-s high temporal-resolution while maintaining noise at a low level. Future work will include a more comprehensive patient study to quantitate the improvement.

**Fig. 10.**
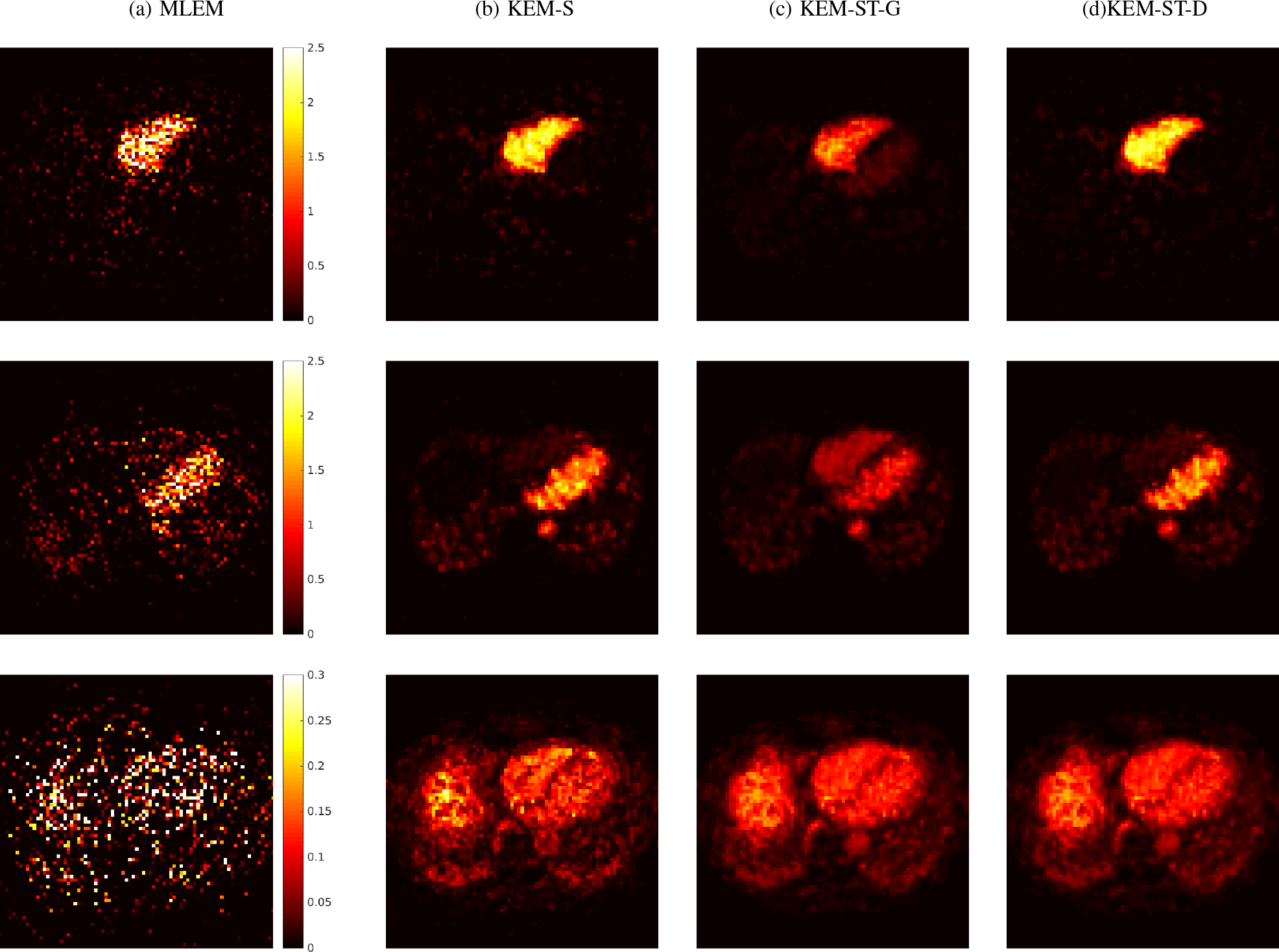
Comparison of different methods for reconstructing HTR patient data at t = 10s (top row), *t* = 20s (middle row) and *t* = 90s (bottom row). (a) MLEM, (b) KEM-S, (c) KEM-ST-G, (d) KEM-ST-D.

**Fig. 11.**
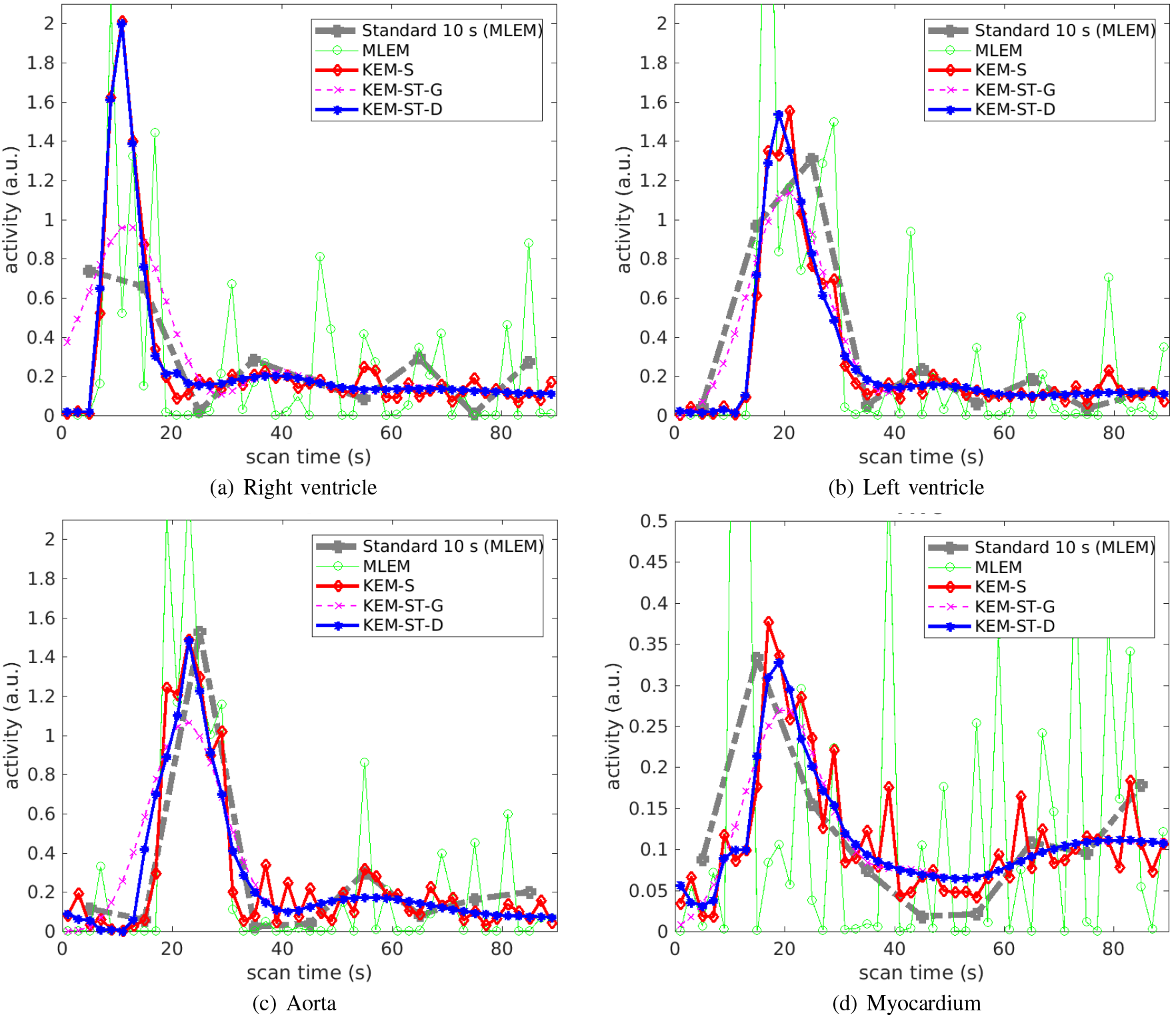
Time activity curves reconstructed by different methods for the pixels in different regions. (a) right ventricle, (b) left ventricle, (c) aorta, (d) myocardium.

## VI. Acknowledgment

The author thanks Dr. Benjamin Spencer and Dr. Yang Zuo for their assistance in patient data acquisition, and Dr. Jinyi Qi for helpful discussion on the topic.

## Notes

This work was supported in part by the NIH R21 HL 131385 and AHA BGIA 25780046. Part of this work was previously presented in the International Meeting on Fully 3D Image Reconstruction in Radiology and Nuclear Medicine, in June 2017.

